# LGR5 and SOX2 expressing progenitor cells in the adult and aged human and mouse inner ear have the potential to produce Myosin 7A positive cells *in vitro*

**DOI:** 10.64898/2026.09.18.752313

**Authors:** Georgina E. Fenton, Tobias Pieper, Cindy Cleypool, Stephanie Sgroi, Francis Rousset, Cecilia de Heus, Nalan Liv, Hans G.X.M. Thomeer, Pascal Senn, Robert J. Stokroos, Louise V. Straatman, Natalia Smith-Cortinez

**Author notes:** Shared first authorship (Georgina E. Fenton, Tobias Pieper).

## Abstract

Hearing loss and deafness as a result of hair cell loss cannot be restored due to the incapacity of spontaneous regeneration of these cells. Compounds that manipulate key signaling pathways can potentially regenerate cochlear hair cells. In order to test different compounds that promote hair cell regeneration *in vitro* models can be used. Three-dimensional cultures have allowed the expansion and experimentation of human and mouse inner ear organoids. This is mainly performed in the embryonic or the early postnatal developmental stage. However, since the majority of patients with hearing loss are adult, it is crucial to understand the adult inner ear regenerative capacity. Here, we evaluated progenitor cell markers in the adult human and mouse inner ear and the generation, expansion and differentiation of cochlear organoids derived from the adult human and mouse inner ear. Cochlear and vestibular sensory epithelium from adult humans; and from adult and aged mice express the progenitor markers SOX2 and LGR5; and can generate organoids *in vitro*. By optimizing the culture conditions, organoids derived from the cochlea and vestibular organ from adult human and adult and aged mouse differentiated to Myosin 7A positive cells. This indicates that the adult inner ear has regenerative capacity. These findings are encouraging for future regenerative therapies to cure hair cell-related hearing loss and deafness.

**Highlights:**

- Human adult cochlear and vestibular organ sensory epithelia possess progenitor potential and express LGR5 and SOX2
- The adult and aged mouse cochlea possesses progenitor potential and express LGR5 and SOX2,
- By optimizing the culture conditions, the adult human and mouse inner ear sensory epithelium can generate organoids *in vitro*
- Human cochlear and vestibular organ derived organoids differentiate to Myosin 7A positive cells with polarized F-Actin.
- Mouse cochlear organoids differentiate to Myosin 7A positive cells with polarized F-Actin, FM1-43 uptake, and stereocilia-like structures

## Introduction

Sensorineural hearing loss in mammals arises from irreversible damage to cochlear sensory cells (hair cells, HCs) and/or their associated neurons (spiral ganglion neurons, SGNs) due to aging, noise exposure or ototoxic medication. According to the World Health Organization, it is expected that more than 900 million people will have disabling hearing loss by 2050 (Wilson et al., 2017). In contrast, hearing loss in non-mammalian vertebrates is reversible as their auditory system possesses regenerative capacity (Brignull et al., 2009). In fish and birds, regeneration of HCs out of supporting cells (SCs) mediates hearing restoration and inducing similar regenerative mechanisms in the mammalian cochlea remains an active area of research. In mammals, SCs in the cochlea give rise to HCs during embryonic development, and these cells express the progenitor cell markers LGR5 and SOX2 (Chai et al., 2012; Chai et al., 2011; Oesterle & Campbell, 2009; Smith-Cortinez, Hendriksen, et al., 2023; Smith-Cortinez, Tan, et al., 2023; Smith-Cortinez et al., 2021; Zak et al., 2015). Although the expression of LGR5 alongside the cochlea greatly diminishes after birth in mice, we (and others) have found LGR5 expression in the third row of Deiters’ cells (DC3), in inner pillar cells and inner border cells up until postnatal day 200 and even after ototoxicity-induced HC loss (Smith-Cortinez, Hendriksen, et al., 2023; Smith-Cortinez et al., 2021). LGR5 positive SCs derived from neonatal mouse cochleae expand and proliferate in 3D cultures in budding-like organoids, hereon called cochlear organoids (Ma et al., 2022; McLean et al., 2017; Xia et al., 2023). These cochlear organoids proliferate under expansion medium containing Wnt agonist and TGFβ inhibitors, and differentiate to HC-like cells after adding differentiation medium containing gamma-secretase inhibitors and opening the epigenetic barrier (Carpena et al., 2025; Ma et al., 2022; McLean et al., 2017; Xia et al., 2023).

Primary cochlear organoid cultures from neonatal mouse cochleae have allowed the generation of drug-testing platforms for evaluation of novel compounds that promote otoprotection and HC regeneration (Chen et al., 2026). Following the work of McLean et al. (2017), optimized culture media have been developed which allow (limited) passaging of neonatal cochlear organoids and a higher yield of HC production (Carpena et al., 2025; McLean et al., 2017; Xia et al., 2023). In addition, neonatal cochlear organoids have been functionalized through co-culture with SGNs which promote HC survival *in vitro* (Carpena et al., 2025; Xia et al., 2023). However, the regenerative capacity of the adult inner ear remains poorly understood. Although LGR5 positive SCs are present in adult mouse cochlea and have been shown to survive after deafening, it is currently unknown whether these cells retain the ability to proliferate or differentiate to HCs *in vitro*.

The expression of LGR5 has not yet been confirmed in the human inner ear. Human progenitor cells have been identified in the vestibular organ (McLean et al., 2017; Senn et al., 2020; van der Valk et al., 2023), which is known to exhibit greater regenerative capacity than the cochlea (Oshima et al., 2007). Isolated cells from the vestibular organ, from adult patients undergoing surgery for vestibular schwannoma, are able to proliferate in culture as inner ear spheroids or organoids and can differentiate to HC-like Myosin 7A positive cells (McLean et al., 2017; Senn et al., 2020). In contrast, the regenerative potential of the human cochlea remains largely unknown, primarily due to the limited availability of adult cochlear tissue. In one study, cochlear progenitor cells isolated from post mortem tissues were shown to proliferate to a very limited extent in culture (Oshima et al., 2007; Senn et al., 2020).

Here, for translational purposes, we evaluated the regenerative capacity of the human adult inner ear within sensory epithelium of the cochlea and vestibular organ from 1) adult patients undergoing surgery for skull base tumors, 2) body donors, and 3) historic temporal bones. We evaluated the expression of progenitor markers like LGR5 and SOX2 in the human cochlea and their potential to proliferate and differentiate in 3D organoids. Furthermore, we evaluated the expansion and differentiation of cochlear organoids derived from adult and aged, Lgr5-GFP (C57BL/6J) transgenic and wild type (WT) C57BL/6J mice in different media compositions.

## Methods

### Human inner ear tissue collection

#### Ex vivo cochlear and vestibular organ sensory epithelium

Adult patients undergoing surgery for skull base tumors (by translabyrinthine and transotic approaches) were included in the study after informed consent. The inner ear tissue collected for this research was obtained according to Dutch legislation and the Code of Conduct for responsibly using human-derived material in health research (www.federa.org). The collection and use of human inner ear tissue was approved by the Biobank Review Committee of the University Medical Center Utrecht (UMC Utrecht) on October 6^th^, 2021 in the protocol titled “Regeneration of human inner ear sensory progenitor cells *in vitro*” (number 21-537). Sensory epithelia— including part of the basal turn of the cochlea and the ampulae and sacculae of the vestibular organ—were collected from patients during the surgery by an experienced surgeon. Age of the patients was recorded anonymously and is presented in **Supplementary Table 1**.

### Fresh post mortem cochlear sensory epithelium

Cochlear sensory epithelium (from basal, middle and apical turns) with a post mortem interval of < 12 hours was collected via a retroauricular transcanal approach (after removal of the otic capsule) from human body donors. Bodies were donated through the body donation program of the Department of Anatomy at the University Medical Center Utrecht, the Netherlands. Informed consent was obtained during life by means of a signed donation form, explicitly permitting the use of these bodies for educational and research purposes. The study was reviewed and approved by the departmental research committee: no additional medical–ethical approval was required. Age of the body donors and post mortem intervals are presented in **Supplementary Table 2**.

### Fixed post mortem cochlear sensory epithelium

Cochlear sensory epithelium (from basal, middle and apical turns) was collected via a retroauricular transcanal approach (after removal of the otic capsule) from temporal bone specimens obtained from the same body donation program of the Department of Anatomy at the University Medical Center Utrecht, the Netherlands. The specimens were obtained from perfusion formaldehyde-fixed bodies whereafter they were stored long term in a preservation fluid containing low concentrations of ethanol, glycerol and phenol.

*Ex vivo* and fresh post mortem human tissue and body donors were transferred to the lab within 30 minutes of collection, in either 1) ice-cold PBS for immunofluorescence microscopy, or 2) ice-cold DMEM/F-12 medium supplemented with B27 and N2 for organoid culture. Before further processing, tissue was inspected and carefully dissected (removing non-sensory epithelial tissue) under a stereomicroscope, on ice.

### Human inner ear tissue dissociation, and culture of organoids

Harvested and dissected human inner ear epithelia from cochlea or maculae were centrifuged and digested in thermolysin (0,5 mg/mL, Sigma-Aldrich) for 20 minutes at 37°C. After washing with PBS and centrifugation, tissues were incubated with Accumax (Innovative Cell Technologies) for 10 minutes at room temperature. After digestion, cells were washed with ice cold PBS, filtered with a 30 µm cell strainer. Red blood cells were lysed with red blood cell lysis buffer (Roche) for 5 minutes. After centrifugation, cells were resuspended in a solution consisting of 60% Matrigel in DMEM/F-12 medium. All isolated cells were plated per well (in three 10 µL drops), in a 24-cell well plate. After the plate was incubated upside-down for 15 minutes at 37 °C (to allow gelation of the Matrigel) 500 µL of isolation medium was added for the first 24h and then changed to expansion medium (EM). The culture medium was replenished every 2-3 days for 15-30 days and cultures were kept in humidified incubators at 37°C and 5% CO_2_. Human inner ear organoids typically become apparent within 7-14 days post-isolation. After the organoids stopped growing, organoids were differentiated in differentiation medium (DM) for 3-10 days.

### Animal models

#### Neonatal mice

C57BL/6J mouse inner-ear tissues were collected under federal animal experimentation license no. 36783 and cantonal license GE395. A total of 20 postnatal mouse pups were used across three independent isolations: eight pups at postnatal day 7 (P7), seven pups at P8 and five pups at P6. Sex was not determined.

### Adult and aged mice

Heterozygous Lgr5-EGFP-IRES-creERT2 mice (Lgr5-GFP; Barker et al. 2007; Stock #008875) were purchased from Jackson Laboratory. In-house breeding was set up to continue the Lgr5-GFP line. In total, 39 Lgr5-GFP and 18 WT mice were used between postnatal day 30-80 (p30-p80, young adult). Further, 6 Lgr5-GFP mice and 13 WT mice aged p100 - p250 (aged). Both males and females were used, and tissue was combined during organoid generation. Detailed experimental animal information is presented in **Supplementary Table S3**. Mice were housed in IVC cages with food and water ad libitum and standard laboratory conditions (12:12 hour light:dark schedule, lights on at 6am). All surgical and experimental procedures were approved by the Dutch central authority for scientific procedures on animals (AVD1150020186105 and AVD11500202417856).

### Genotyping

DNA was extracted from ear tagging tissue to genotype Lgr5-GFP transgenic mice as previously described (Smith-Cortinez et al., 2021). Genomic DNA isolation was performed on 0.2 cm ear clips with DirectPCR lysis reagent (Viagen, Biotech, Los Angeles, CA, USA, 402-E) according to the manufacturer’s instructions. The primers for PCR amplification were: GFP; forward: CACTGCATTCTAGTTGTGG, and reverse CGGTGCCCGCAGCGAG. Amplicons were separated by electrophoresis in a 3% agarose gel.

### Auditory brainstem responses (ABRs)

ABRs were measured as previously described (Smith-Cortinez et al., 2023). Briefly, ABRs were recorded from anaesthetized mice, in a soundproof and electrically shielded box (52 × 34 × 28 cm). We placed 27G subdermal needle electrodes subcutaneous behind the pinna (active), on the skull (reference), and in the hind limb (ground). Acoustic stimuli consisting of 20-μs monophasic clicks were generated and attenuated using a TDT3 system (Multi-I/O processor RZ6; Tucker-Davis Technologies, Alachua, FL, USA) and presented in free field using a Bowers & Wilkins speaker (CCM683; 8 Ω; 25–130 W) at 5 cm distance from the ear. The electrode signals were pre-amplified using a Princeton Applied Research (Oak Ridge, TN, USA) 5113 pre-amplifier (amplification × 5,000; band pass filter 0.1–10 kHz). The amplified signal was digitized by the same TDT3 system for analysis (100 kHz sampling rate, 24-bit sigma-delta converter). Maximum 500 repetitions were averaged and stored for analysis with custom MATLAB software. Attenuation steps of 10 dB were performed, starting with maximum sound level of approximately 105 dB peak equivalent SPL, until 10 dB below the sound level with no visible ABR response.

### Isolation of organ of Corti-derived otic progenitor cells from neonatal mouse

Pups were decapitated, and each head was divided sagittally. The brain and brainstem were removed to expose the temporal bones, which were transferred to a 100-mm Petri dish containing 12 mL ice-cold Hanks’ Balanced Salt Solution (HBSS). The bulla was removed to expose the otic capsule. The bony capsule was then gently separated from the membranous labyrinth. After removal of the modiolus, the cochlear duct was isolated, and the scala media was dissected from base to apex using 5.5 forceps. Dissected cochlear ducts were collected in 1.5-mL microcentrifuge tubes containing ice-cold HBSS. Tissues were dissociated by enzymatic digestion followed by mechanical trituration. Cochlear ducts were incubated in 0.05% trypsin–EDTA (Thermo Fisher Scientific) for 20 min. Enzymatic activity was stopped by adding DMEM/F12 containing HEPES and L-glutamine and supplemented with 1× N2, 1× B27, 1× penicillin–streptomycin and 10% fetal bovine serum (all from Thermo Fisher Scientific). Tissue was mechanically triturated 10–15 times using a P1000 pipette. Cells were collected by centrifugation to remove trypsin and serum; the centrifugation settings were not recorded. The resulting suspension was resuspended in proliferation medium, passed through a 70-µm cell strainer to remove residual undissociated tissue and bone while retaining small organ of Corti-derived cell clusters, and counted. Following centrifugation, cells were adjusted to 100,000 cells in 22.3 µL proliferation medium. All subsequent manipulations before plating were performed on ice. A 60% Matrigel solution (Corning) was prepared in basal medium consisting of DMEM/F12 supplemented with N2 and B27. The cell suspension was mixed with the 60% Matrigel solution at a 1:2 volume ratio. Twenty µL droplets were deposited in the centre of individual wells of a 48-well plate (Greiner), with one droplet per well. Plates were inverted for 15 min to allow gelation, after which 200 µL of the appropriate proliferation or experimental medium was added to each well. On the following day, 100 µL medium was removed from each well and replaced with 200 µL fresh medium. Thereafter, half of the medium was replaced every 3–4 days during a 14-day expansion period. At the end of expansion, the culture medium was replaced with differentiation medium, and half-medium changes were continued every 3–4 days for a further 14 days.

### Adult mouse cochlea and vestibular organ tissue dissociation, and culture of organoids

Mouse inner ears were dissected after CO2 asphyxiation. Under a stereomicroscope cochleae were carefully separated from the vestibular organ and whole cochleae or vestibular organs were crushed in ice cold PBS, centrifuged and digested in thermolysin (0,5 mg/mL Sigma-Aldrich) for 20 minutes at 37°C. After washing with PBS and centrifugation, tissues were incubated with Accumax (Innovative Cell Technologies) for 20 minutes at room temperature. After digestion, cells were washed with ice cold PBS, filtered with a 30 µm cell strainer. Red blood cells were lysed with red blood cell lysis buffer (Roche) for 5 minutes. After counting and centrifugation, cells were resuspended in a solution consisting of 60% Matrigel in DMEM/F-12 medium. Between 100,000 and 150,000 cells were plated per well (in three 10 µL drops), in a 24 well plate. After the plate was incubated upside-down for 15 minutes at 37 °C (to allow gelation of the Matrigel) 500 µL of experimental media was added. The culture medium was replenished every 2-3 days and plates were placed in humidified incubators at 37°C and 5% CO_2_. We evaluated different media compositions. Published media by McLean., et al 2017 (McLean EM and DM) and Xia et al., 2023 (Xia EM and DM) were used, and high growth factor (HGF) medium, Optimized Isolation medium (Opti IM), Optimized Expansion media (Opti1 and Opti2 EM), and Optimized Differentiation media (Opti 1 and Opti 2 DM). Exact media compositions are in **Supplementary Tables S4-9**. Isolation media was used for the first 24h in culture for the protocols from Xia and optimized.

### Adult mouse organ of Corti

Mouse inner ears were dissected after CO_2_ asphyxiation. Under a stereomicroscope cochleae were carefully separated from the vestibular organ. The cochleae were kept in ice cold PBS and the bone was carefully removed from the apical side of the cochlea until the organ of Corti was fully exposed. The organ of Corti was dissected, lateral wall and stria carefully removed and the organ of Corti was processed for the digestion procedure. After dissection, organ of Corti were pooled and centrifuged and digested in thermolysin (0,5 mg/mL Sigma-Aldrich) for 20 minutes at 37°C. After washing with PBS and centrifugation, tissues were incubated with Accumax (Innovative Cell Technologies) for 20 minutes at room temperature. After digestion, cells were washed with ice cold PBS, filtered with a 30 µm cell strainer. Red blood cells were lysed with red blood cell lysis buffer (Roche) for 5 minutes. After counting and centrifugation, cells were resuspended in a solution consisting of 60% Matrigel in DMEM/F-12 medium. Between 100,000 and 150,000 cells were plated per well (in three 10 µL drops), in a 24-cell well plate. After the plate was incubated upside-down for 15 minutes at 37 °C (to allow gelation of the Matrigel) 500 µL of isolation medium was added for the first 24h and then changed to EM for 15 days and in DM for an extra 10 days. The culture medium was replenished every 2-3 days and plates were placed in humidified incubators at 37°C and 5% CO_2_.

### Brightfield microscopy and organoid measurements

Images of the developing organoids were captured using a bright-field inverted microscope (EVOS FL Fluorescence Microscope, catalog number AMF-4305, AMG and/or EVOS XL Core or Agilent BioTek Cytation 5 multimode imaging system.) every 2 to 3 days to keep track of the growth. The appropriate brightfield microscopy objective (e.g., 4x,10x, 20x) was selected to obtain images of the organoids. Organoids were examined to evaluate their sizes, and all discernible (>100 um) organoids were quantified. Organoid area, diameter and circularity were calculated using the ImageJ/Fiji software.

### Cochlear explants

Cochlear explants were harvested from adult C57BL/6J mice (for exact ages and numbers refer to **Supplementary Table 3**). After harvesting, the bone of the cochlea was carefully removed in ice cold PBS and sensory epithelium was dissected, and cultured in ultra-low adherent 24 cell-well plates for 24 hrs in Opti2 isolation medium, 24 hrs in kanamycin (1 mM in basal medium) and 10 days Opti2 DM. After 10 days

### FM1-43 uptake and SirR Actin live staining

Ten or twenty days after isolation, cochlear organoids were incubated for 30 min in SirActin containing the L-type calcium channel blocker verapamil as a broad-spectrum efflux pump inhibitor (as recommended by product manufacturer) in EM or DM. After washing with PBS once, organoids were harvested in ice cold PBS. After centrifugation, organoids were incubated for 30 sec or 1 min in ice cold FM1-43fx (5 µg/mL in DMEM/F-12 medium), followed immediately by the addition of 4% paraformaldehyde (PFA) and then centrifuged. After carefully removing the supernatant, organoids were incubated in 4% PFA overnight at 4C. After fixation, samples were processed for immunofluorescence microscopy.

### Immunofluorescence microscopy

Immunofluorescence staining was conducted on human and mouse whole cochlear turns or whole organoids. The organoids were harvested by collecting them in cell recovery solution (using wide-tipped 1000 µL FCS-coated pipette tips). After centrifugation and washing with PBS 3 times, samples were in PFA overnight at 4°C and permeabilized (0.5% Triton X-100). Later, samples were incubated with a blocking solution (2% donkey serum and 0.1% Triton X-100 in PBS) for 1h at room temperature or overnight at 4°C. Later, organoids were incubated with primary antibodies, anti-myosin VIIA (MYO7A, 1/200, rabbit, Proteus Biosciences, 25-6790), anti-Ki-67 (1/200, rat, Thermo), anti-GFP (1/500, goat, Abcam, ab5450), or anti-LGR5 (1/200, rabbit, proteus) during an overnight incubation at 4°C. Afterwards, samples were washed 3 times with blocking solution and were then subjected to incubation with secondary antibodies and/or DAPI and/or phalloidin-rhodamine for 90 minutes at room temperature (donkey-anti Rabbit-Alexa 594 (1/500, Invitrogen, A-21207), donkey-anti Rat 594 (1/500, A48271, Invitrogen), donkey-anti Goat-Alexa 488 (1/500, Abcam, AB150129), and DAPI solution (1/500, Abcam, AB228549). Lastly, the specimens were washed in PBS, cleared using a fructose-glycerol solution and mounted using Vectashield Antifade Mounting Medium (Vector laboratories, H-1000). Slides were imaged with a Zeiss LSM700 Confocal Microscope or a Zeiss LSM800 confocal microscope. Three-dimensional image reconstruction of Z-stacks, as well as the analysis, were performed using the ImageJ/Fiji software.

### Ultrastructural Analysis with Electron Microscopy

Whole Matrigel drops were washed twice with warm PBS (37C) and fixed in half-strength Karnovsky’s fixative (2.5% glutaraldehyde [EMS] and 2% formaldehyde [Sigma]) prepared in 0.1 M PHEM buffer (pH 7.4) at room temperature for 2 hours. After fixation, the samples were rinsed thoroughly with PHEM buffer and stored in 1% formaldehyde at 4°C until further processing. Post-fixation was performed in 1% osmium tetroxide (OsO₄) and 1.5% potassium ferricyanide (K₃Fe(CN)₆) in 1 M phosphate buffer (pH 7.4) for 2 hours at room temperature. Samples were then dehydrated through a graded acetone series (70% overnight, followed by 90% for 15 min, 96% for 15 min, and 100% acetone for 3 × 30 min) and embedded in Epon resin (SERVA). Ultrathin sections (70 nm) were cut using a Leica Ultracut UCT ultramicrotome, mounted on formvar- and carbon-coated TEM grids, and stained with uranyl acetate and lead citrate using a Leica AC20 staining system.

Electron micrographs were acquired using a JEOL JEM1011 transmission electron microscope equipped with a Veleta 2k × 2k CCD camera (EMSIS, Münster, Germany). Image visualization and analysis were conducted in Fiji (Schindelin et al., 2012. Nature Methods, 9(7):676–682. doi:10.1038/nmeth.2019).

### Statistical analysis

Data analysis was executed using GraphPad Prism 10. Mean values with corresponding standard deviations (SD) are presented (refer to Figure Legends for additional details). Statistical significance was determined through one-way ANOVA with Tukey’s multiple comparison test or unpaired student t test where appropriate. Significance levels are denoted by asterisks: P ≤ 0.05 (∗), P ≤ 0.01 (∗∗), or P ≤ 0.001 (∗∗∗) or P < 0.0001 (∗∗∗∗). Details specific to each figure are provided in the corresponding figure legends.

## Results

### Expression of progenitor cell markers in the adult and aged human and mouse cochlea

Cochlear sensory epithelium was collected from fixed post mortem, fresh post mortem and *ex vivo* samples. Our surgical technique allowed us to extract the whole basal, middle and apical turns in fixed and fresh post mortem samples, and only a small biopsy from the basal turn of the cochlea in *ex vivo* live donors; all of which after dissection preserved the organ of Corti intact (brightfield images, **Figure 1A**). In fixed and fresh post mortem organ of Corti, we observed Myosin 7A (in red, **Figure 1A**) and F-actin expression (in white, **Figure 1A**) in inner and outer HCs; We observed prestin expression in outer HCs (in orange, top and middle panels, **Figure 1A**). Importantly, we observed a highly preserved anatomical structure of stereocilia and Prestin in samples derived from fresh post mortem (middle panels, **Figure 1A)**. In *ex vivo* samples, Myosin 7A expression was limited (in red, bottom panels, **Figure 1A**), most likely because vestibular schwannoma patients have in general severely reduced hearing thresholds due to the tumors. Immunostaining of key progenitor cell markers showed SOX2 expression in all SCs of the organ of Corti in fixed and fresh post mortem samples (in purple, top and middle panels, **Figure 1A**; and expression of LGR5 in the organ of Corti (F-Actin positive) of fixed post mortem samples (LGR5, green, F-Actin (white) top panels, **Figure 1A**).

**Figure 1:**
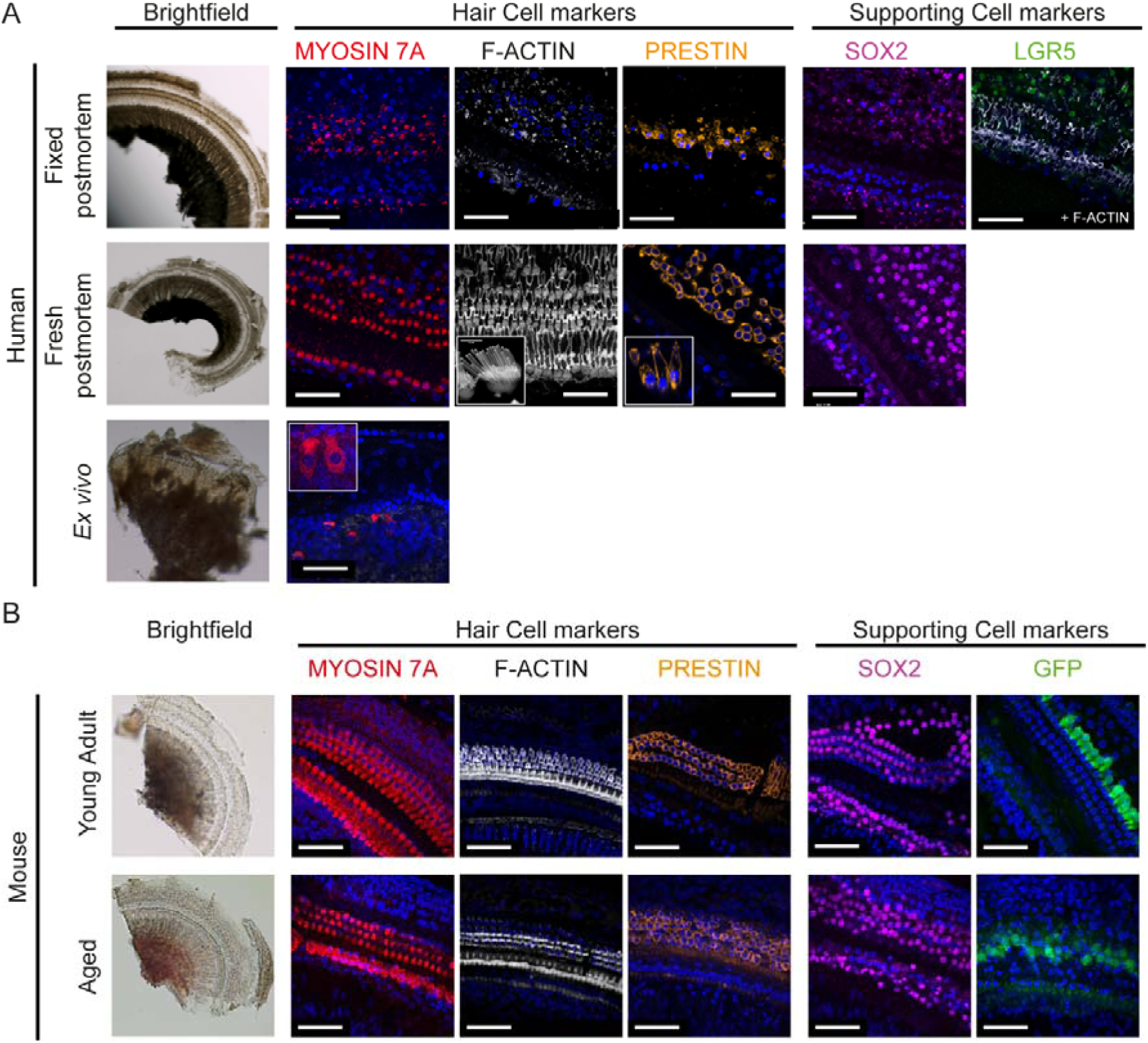
Characterization of progenitor cells found in the young adult and aged human and mouse organ of Corti. A) Adult and aged human organ of Corti was obtained and dissected from fixed post mortem, fresh post mortem and *ex vivo* cochlear tissue. Samples were obtained after a translabyrinth approach to expose the organ of Corti and the full labyrinth was removed from fixed and fresh post mortem samples; and a small biopsy was taken from the *ex vivo* sample. Representative images of bright field microscopy, and immunofluorescence microscopy for the hair cell markers Myosin 7A (red), F-Actin (white), Prestin (orange), and progenitor cell markers SOX2 (purple) and LGR5 (green). B) Cochleae were harvested from young adult (p30 – p80) and aged (p100 - p250) C57BL/6J and Lgr5-GFP transgenic mice. After fixation, cochleae were dissected and the organ of Corti was processed for immunostaining for the hair cell markers Myosin 7A (red), F-Actin (white), Prestin (orange), and progenitor cell markers SOX2 (purple) and GFP (LGR5, green).

To investigate the changes of progenitor cell markers during ageing, cochlear sensory epithelium was collected from young adult and aged transgenic Lgr5GFP (C57BL/6J background) mice. After fixation, the organ of Corti was preserved intact (brightfield images, **Figure 1B**). Immunostaining of key HC markers showed expression of Myosin 7A (in red, top panels, **Figure 1B**) and F-Actin (in white, top panels, **Figure 1B**) in inner and outer HCs and Prestin (in orange, top panels, **Figure 1B**) in outer HCs. In aged mouse organ of Corti, there was reduced Myosin 7A expression (in red, bottom panels, **Figure 1A**), F-Actin expression (in white, bottom panels, **Figure 1B**); and Prestin expression (in orange, bottom panels, **Figure 1B**) indicative of early onset of hearing loss. Indeed, ABR thresholds and wave I amplitudes were significantly reduced in aged compared to young adult mice (**Supplementary Figure 1** and **Supplementary Table 10**). Immunostaining of key progenitor cell markers showed SOX2 expression (in purple, bottom panels, **Figure 1B**) in all SCs in the organ of Corti and expression of GFP (LGR5) in young adult and aged mice (in green, bottom panels, **Figure 1B)**.

In conclusion, these results suggest the existence of SOX2 positive and LGR5 positive progenitor cells in the adult and aged human and mouse organ of Corti.

### Adult Mouse and Human Inner Ear Progenitors Produce Organoids

To evaluate proliferation and differentiation capacity of the human adult inner ear, we promoted growth and proliferation of *ex vivo* human inner ear epithelium collected from patients (see **Supplementary Table 1** for patient characteristics). The mean was 49.3 years and the median 51 years, so most patients were mature adults. Cochlea or vestibular organ epithelium was digested to single cells and cultured in McLean EM [McLean et al 2017]. After 3 days in culture, digested single cells were able to form clusters of cells (organoids) from variable sizes up to 20 days (**Figure 2A)**. Organoids were seen in 6/10 patients in cochlear-derived epithelium, and 10/10 patients in vestibular organ-derived epithelium (**Table 1**). The number of organoids was significantly larger in samples derived from vestibular organ epithelium compared to cochlear epithelium (**Figure 2B** and **Table 2**). Cochlear and vestibular organ-derived organoids were able to produce a limited number of Myosin 7A positive cells after 3 days of differentiation (**Figure 2C**). These results suggest the adult and aged human inner ear has (limited) regenerative capacity *in vitro*.

**Figure 2.**
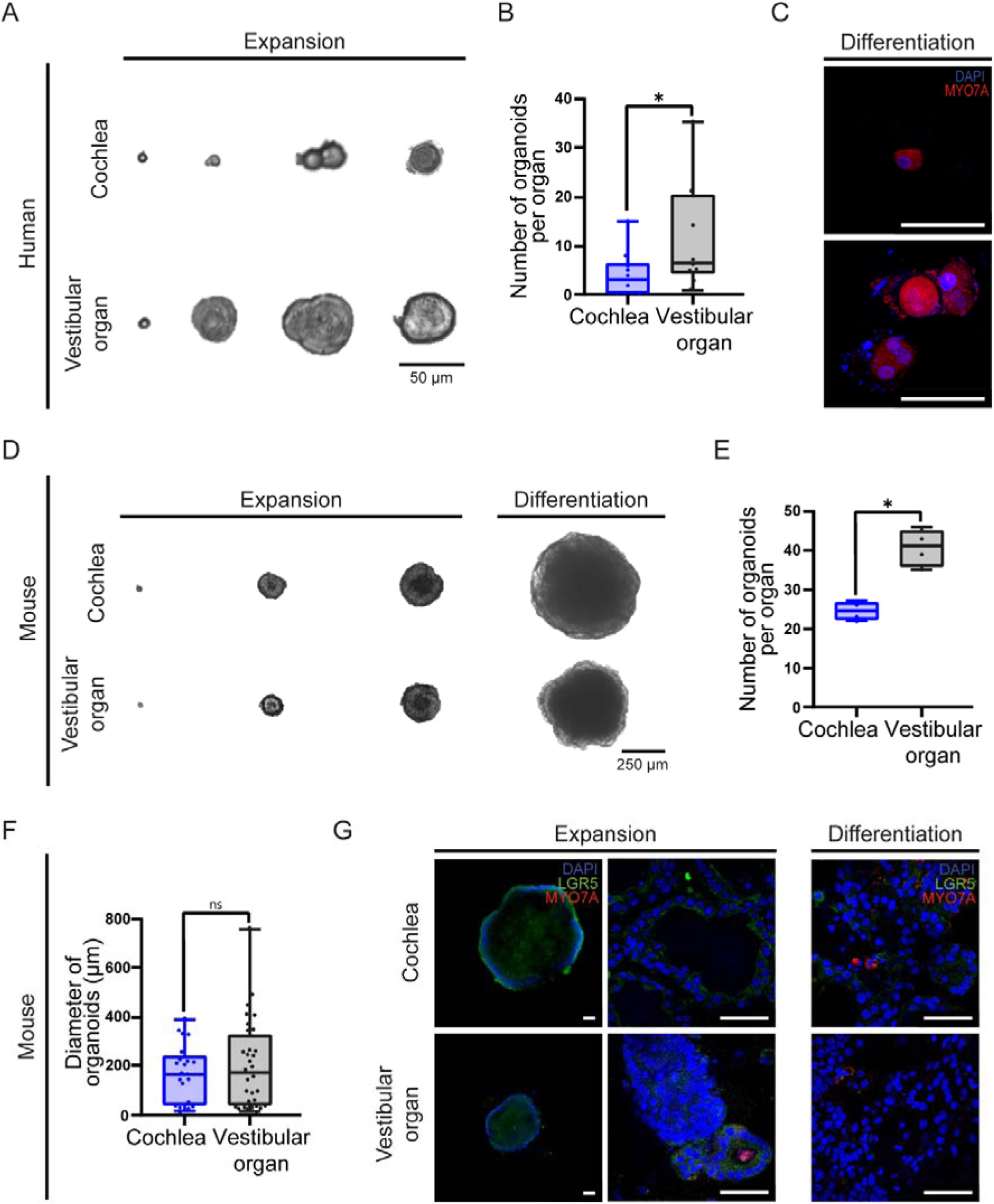
Human and mouse inner ear organoids grow in McLean medium. A-C) Progenitor cells from the inner ear (cochlea and vestibular organ) of 10 adult patients (samples from patients #1-10) were grown in culture in McLean 2017 expansion medium (EM) over 15 days and A) representative images were taken by brightfield microscopy on days 3, 6, 10 and 15. B). Representative images of immunofluorescence microscopy of the hair cell marker (Myosin 7A, in red) in organoids derived from cochlea and vestibular organ from patients. n=3. C) Number of organoids before the start of differentiation. D-G) Progenitor cells from the inner ear (cochlea and vestibular organ) of adult Lgr5-GFP mice (5 Lgr5-GFP mice aged p250) were grown in culture in McLean 2017 (EM) over 10 days and D) Representative images were taken by brightfield microscopy on days 3, 6, 10 and 20. E) Number of organoids per organ at day 10. F) Diameter of organoids in micrometers derived from cochlea and vestibular organ from aged Lgr5-GFP mice. G) Representative images of immunofluorescence microscopy of the progenitor cell marker, Lgr5-GFP, and the hair cell marker, Myosin 7A, in organoids derived from cochlea and vestibular organ from aged Lgr5-GFP mice.

To develop a translational model for cochlear regeneration in ageing, we evaluated the proliferation and differentiation capacity of LGR5 positive SCs derived from cochlea and vestibular organ of aged (>p250) Lgr5-GFP transgenic mice and generated 3D cochlear- and vestibular organ-organoid cultures in McLean EM [McLean et al., 2017]. On the isolation day, cultures consisted of single cells, organoids were visible 3 - 4 days after isolation and continued to grow the first 10 days in expansion conditions (**Figure 2D**). On day 10, we observed significantly more organoids from vestibular organ than from cochlea (**Figure 2E**). There were no differences in the size of the organoids (diameter) between organoids derived from cochlea and vestibular organ (**Figure 2F**). In expansion conditions, organoids derived from cochlea and vestibular organ expressed a limited number of LGR5 positive cells (**Figure 2G**). After differentiation, limited to no Myosin 7A positive cells were observed in both cochlea- and vestibular organ-derived organoids (**Figure 2G**). These experiments suggest the aged inner ear has LGR5 positive cells which have the capacity to proliferate; however, the differentiation capacity in the published media used is limited.

### Optimized Medium Enhances Proliferation and Differentiation of LGR5 positive Progenitors in Adult Mouse Cochlear Organoids

To increase proliferation and differentiation capacity of LGR5 positive SCs, different published and optimized (**Supplementary Table S4-9**) mediums were compared for the culture of cochlear organoids derived from young adult mice. Expansion medium McLean EM [McLean et al., 2017] was compared to Xia EM [Xia et al., 2023], and optimized (previously unpublished) media high growth factor (HGF), Optimized 1 (Opti1) and Optimized 2 (Opti2) EMs. On the day of the isolation the cultures derived from young adult (p30-p60) Lgr5-GFP mice consisted of single cells, cochlear organoids were visible 3-4 days after isolation in all mediums tested. On day 10, we observed an average of 10 organoids per cochlea in McLean EM, 8 organoids per cochlea in Xia EM, 20 organoids per cochlea in HGF EM and Opti1 EM and 25 organoids per cochlea in Opti2 EM (**Figure 3A**). The organoids had highly variable sizes, also within same EM (**Figure 3B**). At day 10, some organoids were harvested and some were placed in DM for 10 days. In expansion conditions, organoids in McLean EM, Xia EM and HGF EM expressed one or two LGR5 positive cells per organoid (**Figure 3C**). In contrast, organoids grown in Opti1 EM and Opti2 EM had significantly more LGR5 positive cells than the other media (**Figure 3C**). The LGR5 positive cells were located in a structured row of cells at the rim of the organoids grown in Opti2 (**Figure 3C**). In differentiation conditions, only one or two Myosin 7A positive cells were observed in McLean DM, Xia DM or in HGF DM (**Figure 3C**). In Opti1 DM and Opti2 DM, enhanced Myosin 7A expression was observed compared to the other media (**Figure 3C**). In Opti2 DM, Myosin 7A positive cells were found in the rim of the organoids and had a pear-like shape (**Figure 3C**). Interestingly, larger organoids showed less Myosin 7A expression in Opti2 DM (**Supplementary Figure 2**), suggesting the size of the organoid impacts its ability to differentiate into Myosin 7A positive cells.

**Figure 3:**
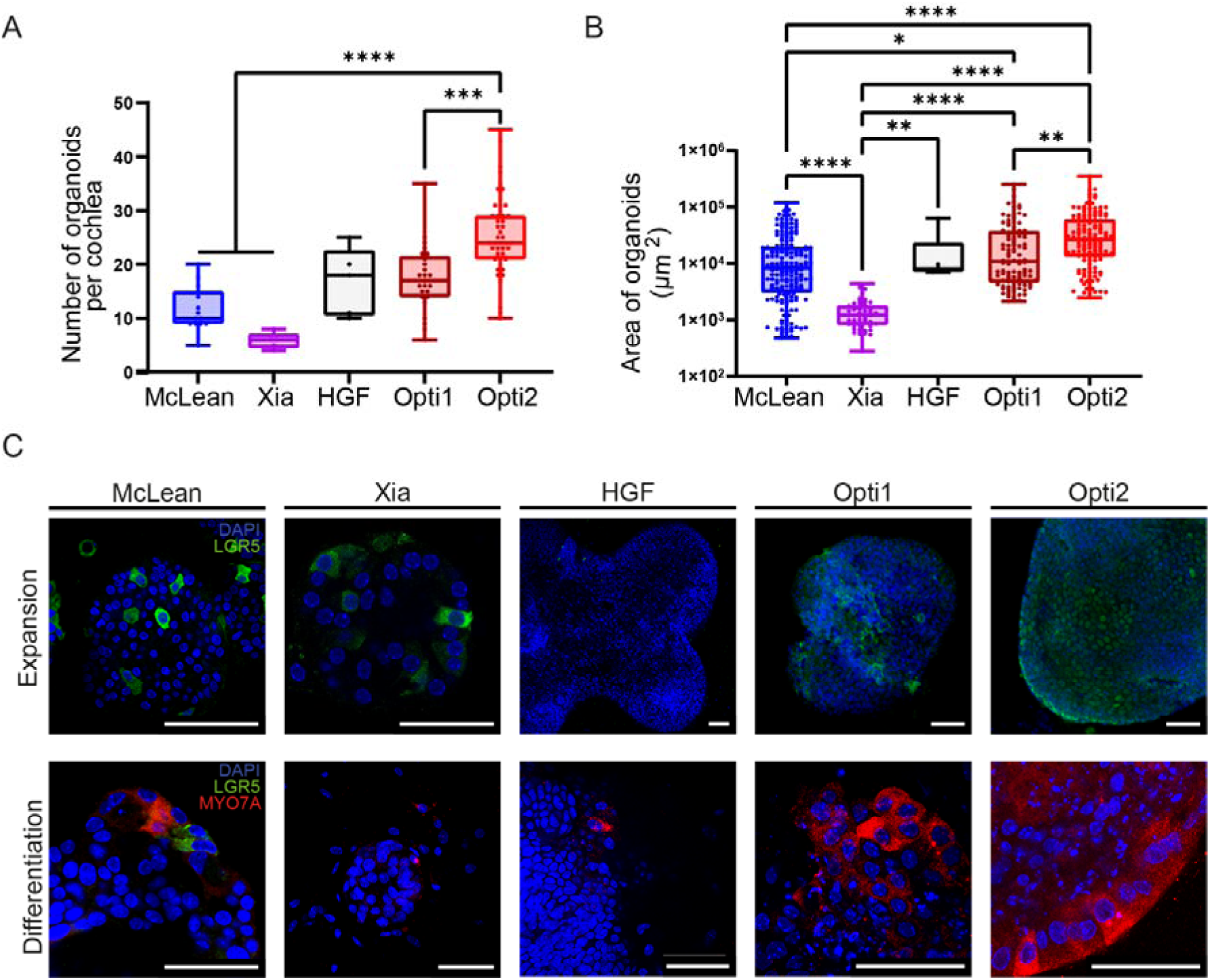
Growth, expansion and differentiation of mouse cochlear organoids in optimized conditions. A) Number of cochlear organoids per cochlea present at day 10 in different expansion media (EM) tested, McLean EM, Xia EM, High growth factor (HGF), Opti1 and Opti2. (Kruskal-Wallis test. H(4) = 48.26, P < 0.0001). B) Area of cochlear organoids in micrometers present at day 10 in different expansion media (EM) tested, McLean EM, Xia EM, High growth factor (HGF), Opti1 and Opti2. (Kruskal-Wallis test. H(4) = 192.7, P < 0.0001. n = 31 Mclean organoids, n = 54 Xia organoids, n = 6 HGF organoids, n= 109 Opti1, n = 140 Opti2). C) Representative images of immunofluorescence microscopy of the progenitor cell marker (LGR5-GFP, in green) and the hair cell marker (Myosin 7A, in red) in cochlear organoids derived from Lgr5-GFP adult mice in expansion (top panels) and differentiation (bottom panels) media tested (McLean EM, Xia EM, High growth factor (HGF), Opti1 and Opti2). Scale bar=50 um.

In conclusion, cochlear organoids derived from adult mice grow and proliferate in culture using different media compositions; however limited LGR5 expression is observed. More importantly, limited Myosin 7A expression was observed in differentiation conditions using the published media for neonatal cochlear organoids. Using an optimized medium (Opti2) we observed enhanced LGR5 expression at the rim of the organoids under expansion conditions and enhanced Myosin 7A expression in pear-shaped cells located at the rim of the organoids in differentiated conditions.

### Adult and Aged Mouse Cochlear Organoids Produce Myosin 7A positive HC-like Cells in Optimized Medium

To characterize the cochlear organoids produced in the optimized condition (Opti2 EM), we evaluated the growth of organoids from the cochlea of young adult (p30-60) and aged (>p250) WT and Lgr5-GFP mice. Young adult WT and Lgr5-GFP mice had normal hearing ABR thresholds (**Supplementary Figure 1A** and **Supplementary Table S10**) while aged WT and Lgr5-GFP mice had reduced ABR thresholds and reduced amplitudes in wave I, suggestive of aged-induced hearing loss (**Supplementary Figure 1B** and **Supplementary Table S10**). The number of organoids generated per cochlea was similar in young adult WT versus young adult Lgr5-GFP mice; however, it was significantly reduced in aged Lgr5-GFP mice compared to aged WT mice (**Figure 4A**). The size of the organoids was significantly smaller in young adult Lgr5-GFP mice compared to young adult WT mice (**Figure 4B**) and in aged WT compared to young adult WT (**Figure 4B**).

**Figure 4:**
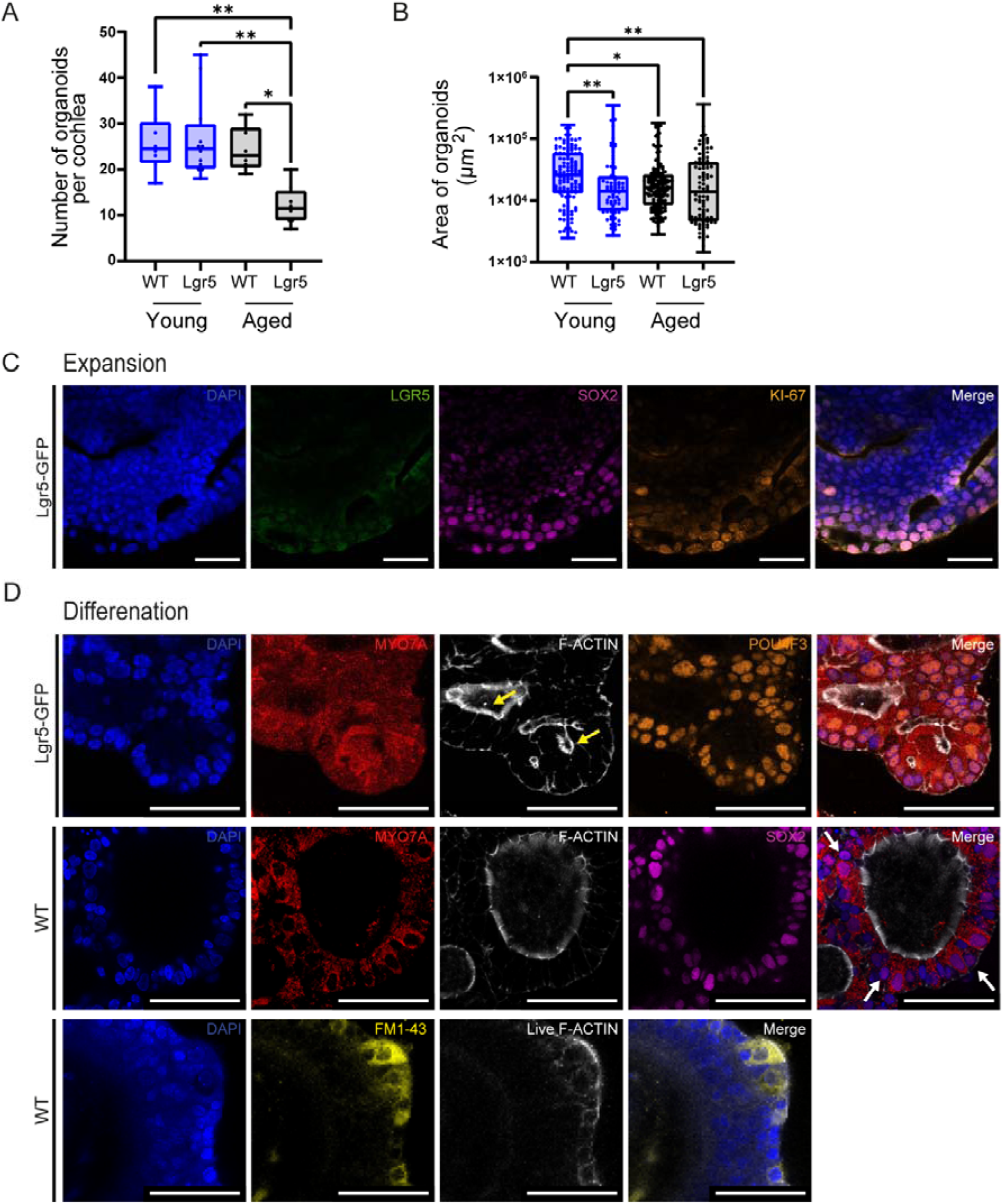
Characterization of mouse cochlear organoids in Opti2 Expansion and Differentiation Media. A) Number of cochlear organoids per cochlea present at day 10 in Opti2 expansion medium (EM) (Mann-Whitney test. U = 188.5, *P* = 0.0162). B) Size of cochlear organoids in micrometers present at day 10 in Opti2 EM (Mann-Whitney test. U = 8570, *P* = 0.0003. n = 140 young organoids measured, n = 162 aged organoids measured). C) Representative images of immunofluorescence microscopy of progenitor cell markers (Lgr5-GFP, SOX2, Ki67) in cochlear organoids derived from Lgr5-GFP and WT adult mice in Opti2 EM. D) Representative images of immunofluorescence microscopy of progenitor cell markers (Lgr5-GFP, SOX2, Ki67) and hair cell markers (Myosin 7A, F-Actin, POU4F3, FM1-43) in cochlear organoids derived from Lgr5-GFP and WT adult mice in Opti2 differentiation medium (DM).

After differentiation, we evaluated the expression of the progenitor cell markers (LGR5, SOX2) the proliferation marker (Ki67) and the HC markers (Myosin 7A, F-Actin, POU4F3), and performed uptake of FM1-43 in the presence of SiR Actin (F-Actin probe). In Opti2 EM, organoids expressed Ki67 positive nuclear staining (**Figure 1C**, in orange). Organoids showed SOX2 positive nuclear staining (**Figure 1C**, in purple) and expressed LGR5-GFP in Lgr5-GFP mouse derived organoids (**Figure 1C**, in green). F-Actin (**Supplementary Figure 2**, in white) had low expression throughout the organoids; there was no specific Myosin 7A expression (**Supplementary Figure 3**, in red); nor uptake of FM1-43 (**Supplementary Figure 3**, in yellow). After differentiation, cells within organoids expressed robust Myosin 7A in cytoplasm with enrichment in the apical side (**Figure 4D**, in red); organoids had nuclear POU4F3 expression (**Figure 4D**, in orange) and nuclear SOX2 expression (**Figure 4D**, in purple). SOX2 staining was present mostly in Myosin 7A positive cells but also in Myosin 7A negative neighboring cells (**Figure 4C**, white arrows). F-Actin (**Figure 4D**, in white) was polarized and enriched in the apical side of Myosin 7A positive cells; and in some Myosin 7A positive cells F-Actin was expressed in stereocilia-like structures (**Figure D**, yellow arrows). FM1-43 uptake was specifically in cells that showed polarized and enriched F-Actin in stereocilia-like structures (**Figure 4D**, in yellow). Similar results were obtained for differentiation of cochlear organoids from young adult and aged WT and Lgr5-GFP mice. Electron microscopy analysis of differentiated cochlear organoids revealed the presence of highly polarized cells with clear cuticular plates (**Figure 5A**), tight junctions between differentiated cells with presence of elongated ciliated structures in the apical side (**Figure 5B**), which contained microtubules characteristic of kinocilium (**Figure 5C**) and stereocilia-like actin arrangements (**Figure 5D**).

**Figure 5:**
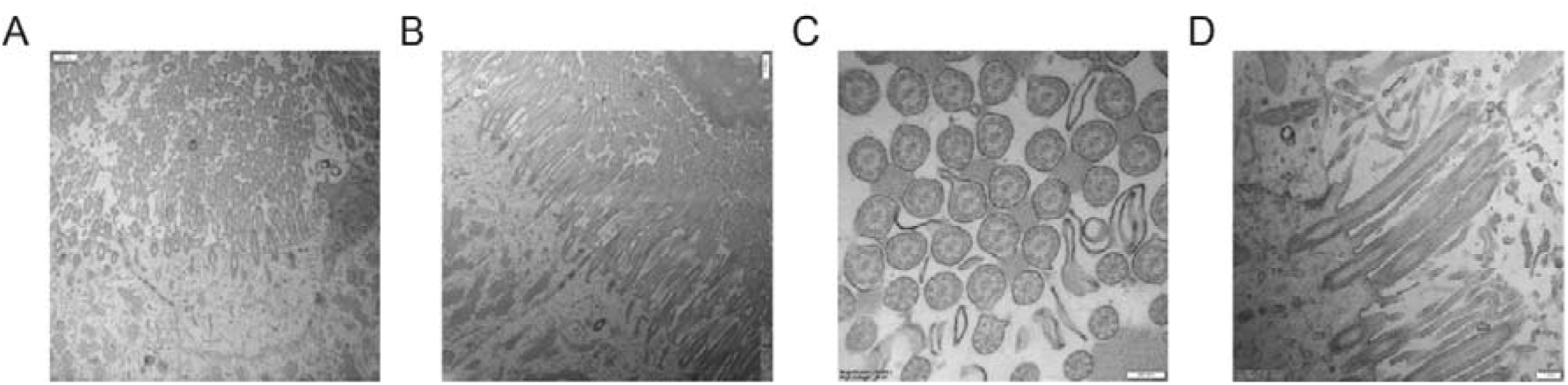
Electron microscopy of cochlear organoids in Opti2 Differentiation media. Representative images of electron microscopy of cochlear organoids derived from adult WT mice in Opti2 differentiation medium (DM) showing A) Apical side of a differentiated cell; B) tight junctions between differentiated cells and ciliated structures in the apical side; C) example of microtubule-like structures present in the ciliated structures; and D) example of stereocilia-like actin-based structures in the apical side of differentiated cells.

Overall, these results suggest that the cochlear organoids derived from adult mice can differentiate to Myosin 7A positive HC-like cells using Opti2 DM.

### Neonatal and adult Mouse organ of Corti produce Myosin 7A positive HC-like Cells In vitro

We dissected the organ of Corti of adult and neonatal mouse cochleae, made single cell suspensions and evaluated organoid growth 14 days after culture initiation. The number and size of organoids was significantly larger in samples derived from the neonatal mouse organ of Corti than from the adult mouse organ of Corti (**Figure 6A**). After differentiation, cells within organoids expressed robust Myosin 7A expression (in red), polarized F-Actin (in white) and nuclear POU4F3 expression (in purple) in organoids derived from adult (top panels, **Figure 6B**) and neonatal (middle panels, **Figure 6B**) WT mice. Neonatal mouse cochlear organoids had uptake of FM1-43 (bottom panels, **Figure 6B**). These results confirm that progenitor cells from the neonatal and adult mice organ of Corti can differentiate to Myosin 7A positive cells *in vitro*.

**Figure 6.**
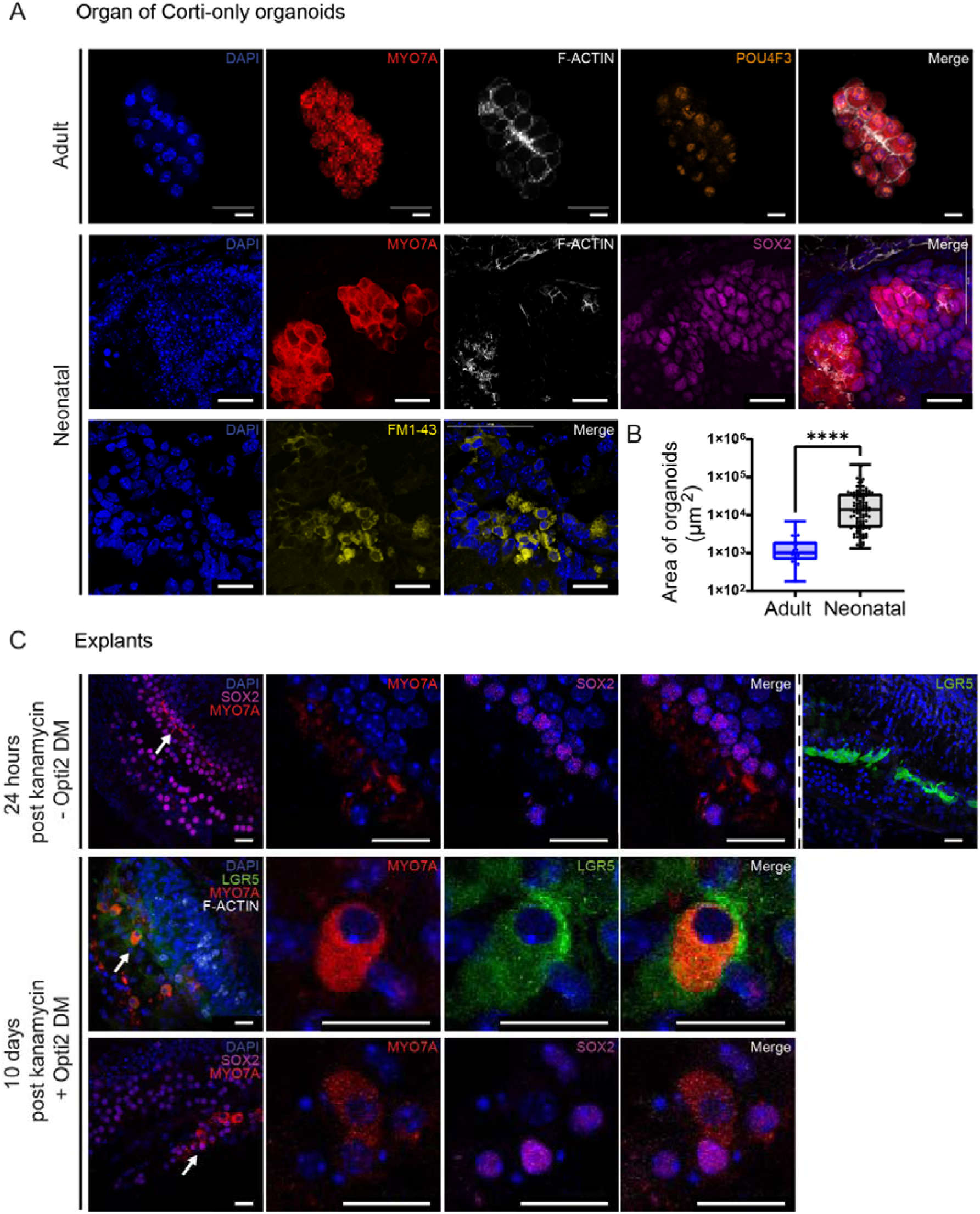
Opti2 DM induces the differentiation of newly produced Myosin 7A from adult organ of Corti *in vitro* and *ex vivo*. Cells from the cochlear sensory epithelium of neonatal and adult WT mice were grown in culture in Opti2 expansion medium (EM) for 14 days. A) Size of organoids in micrometers, B) Representative images of immunofluorescence microscopy of progenitor cell markers (Lgr5-GFP, SOX2, Ki67) and hair cell markers (Myosin 7A, F-Actin, POU4F3, FM1-43) in cochlear organoids derived from adult and neonatal WT mice in Opti2 differentiation medium (DM). C) Representative images of immunofluorescence microscopy of progenitor cell markers (Lgr5-GFP, SOX2, Ki67) and the hair cell marker (Myosin 7A) in cochlear explants isolated from adult WT mice, cultured for 24 hrs in isolation medium, exposed to kanamycin (1mM) for the following 24 hrs and cultured for the following 10 days in Opti2DM.

### Cochlear Explants Produce Myosin 7A positive Cells In Situ

Cochlear explants were exposed to kanamycin (1 mM, 24h) in culture 1 day after isolation to induce HC death and cultured for up to 10 days in Opti2 DM. One day after Kanamycin, organ of Corti expressed SOX2 (in purple) in all SCs, LGR5 (GFP, in green) in DC3s and limited Myosin 7A (in red, top panels, **Figure 6C**). Ten days after kanamycin, in Opti2 DM culture, organ of Corti showed expression of Ki67 (in white) suggestive of proliferation induction; there was newly produced Myosin 7A positive cells (in red) colocalizing with LGR5 (GFP, in green, middle panels, **Figure 6C**). SOX2 (in purple) positive cells were present in the vicinity of the newly produced Myosin 7A positive cells (bottom panels, **Figure 6C**). These results show that the Opti2 DM is able to induce proliferation and differentiation of new Myosin 7A positive HC-like cells in situ in mouse adult-derived cochlear explants.

### Human Organ of Corti Produce Myosin 7A positive HC-like Cells In vitro

We dissected the organ of Corti of adult and neonatal mouse and from *ex vivo* human inner ear sensory epithelia (organ of Corti and maculae). *Ex vivo* human organ of Corti and maculae sensory epithelium were able to produce organoids (**Figure 7A&B**). Macculae-derived organoids (n=46) were significantly bigger than organ of Corti-derived organoids (n=12, **Figure 5B**). After differentiation, cells within organoids expressed robust Myosin 7A expression and polarized F-Actin in all organoids assessed (human organ of Corti, human maculae-derived organoids (**Figure 7C**). These results confirm that progenitor cells from the human inner ear are able to differentiate to Myosin 7A positive cells *in vitro*.

**Figure 7:**
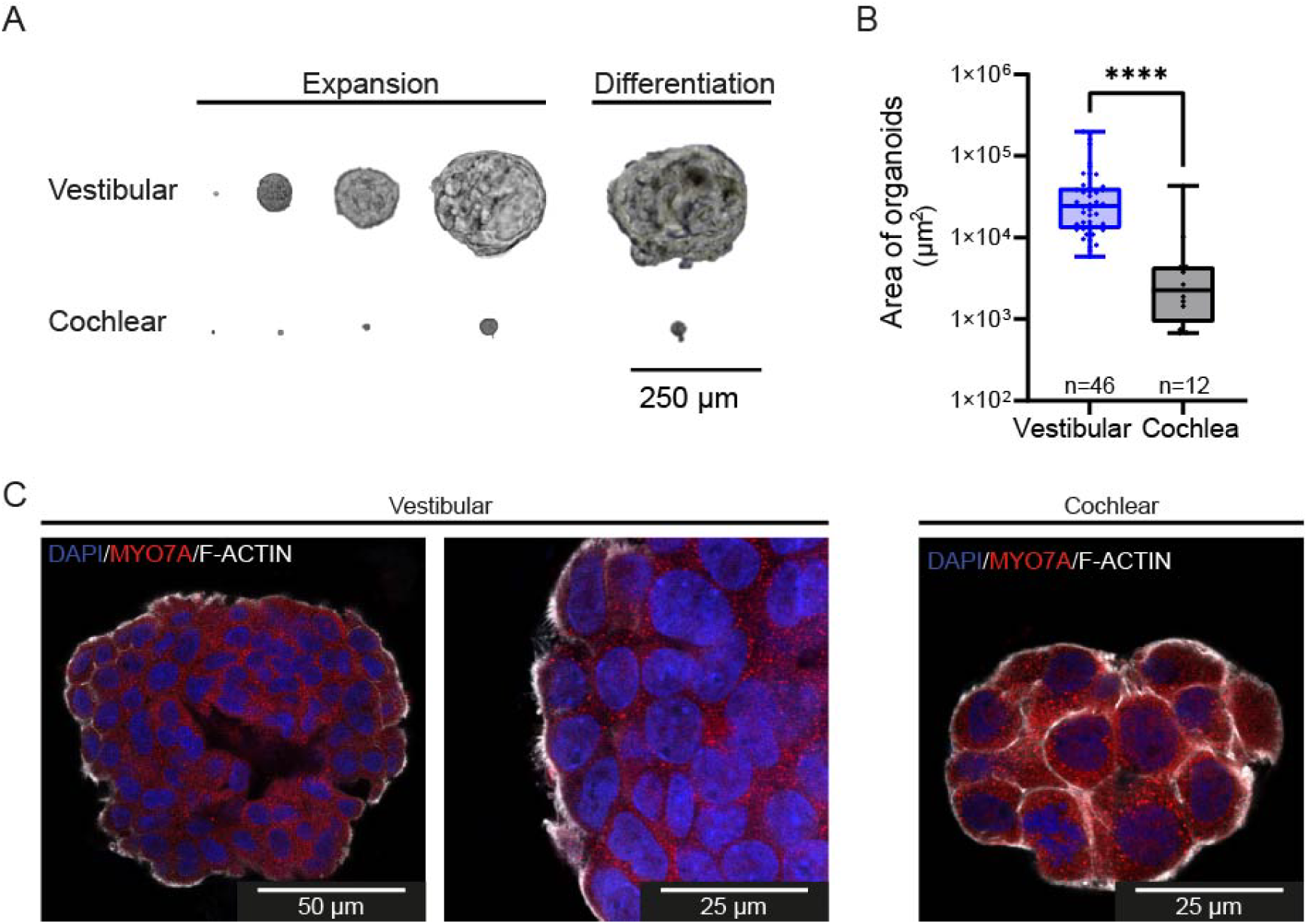
Human inner ear sensory epithelium produces Myosin 7A positive cells *in vitro*. Progenitor cells from the sensory inner ear epithelium (organ of Corti and maculae) of 1 adult patient (aged 23 years old) grown in culture in Opti2 expansion medium (EM) for 15 days and in differentiation medium (DM) for 10 days. A) Representative images were taken by brightfield microscopy on days 1, 6, 10, 15 and 25. B) Number of organoids per organ before start of differentiation (Mann-Whitney test. U = 42, *P* < 0.0001. n = 46 vestibular, n = 12 cochlear). C) Representative images of immunofluorescence microscopy of the hair cell marker Myosin 7A (red) and F-Actin (white,) and DAPI (blue).

## Summary

Here, we evaluated the regenerative potential of the adult and aged mouse cochlea and the adult and aged human inner ear. We histologically characterized the human organ of Corti harvested within 12 hrs post mortem from body donors. We evaluated the expression of progenitor cell (LGR5 and SOX2) and hair cell markers (Myosin 7A, Prestin, F-actin) and showed for the first time that LGR5 and SOX2 progenitor cell markers are expressed in the human adult cochlea, even at an advanced age. Additionally, we evaluated the regenerative capacity of the human organ of Corti by culturing isolated cells from discarded surgical tissues from tumor resection surgeries from vestibular schwannoma patients. Culture of cochlear organoids from these samples demonstrated that by using an optimized protocol, human cochlear and vestibular organ cells can proliferate in culture and can differentiate to Myosin 7A positive/F-Actin positive cells *in vitro*.

Further, we evaluated the capacity of cochlear (and vestibular organ) cells derived from adult and aged Lgr5-GFP and WT mouse to produce cochlear organoids and to differentiate to HCs *in vitro*. We optimized the culture conditions for adult cochlear progenitor cells; when using published media for neonatal cochlear organoids (Carpena et al., 2025; McLean et al., 2017; Xia et al., 2023) in adult derived progenitor cells, organoids were generated. However, these published media promoted limited LGR5 expression in the proliferation phase and only a few Myosin 7A positive cells were observed in the differentiation phase. With the use of surrogate Wnt, as a water-soluble and highly potent Wnt agonist, and R-spondin-1 (Rspo1) in the isolation and proliferation phase of the cochlear organoid generation, we observed enhanced proliferation and LGR5 expression. With the use of sonic hedgehog activation, Notch inhibition and a reduced Wnt signaling activation (addition of CHIR99021, removal of Rspo1 and Surrogate Wnt) we promoted the differentiation to Myosin 7A positive and F-Actin positive cells. Using the optimized medium, differentiated cells showed expression of POU4F3 and SOX2. Upon maturation of these cells, they showed elongated F-Actin in stereocilia-like structures, confirmed by electron microscopy and incorporated the neural probe FM1-43, suggesting the presence of functional mechanotransduction (MET) channels. This was replicated in cochlear progenitor cells derived from dissected neonatal and adult mouse organ of Corti.

We evaluated the capacity of the optimized medium to promote proliferation and differentiation of cochlear progenitor cells in situ in adult cochlear explants in an *in vitro* ototoxicity model. We showed cochlear progenitor cells were able to proliferate and regenerate new Myosin 7A positive cells after kanamycin induced damage.

These results strongly suggest that progenitor cells are present in adult patients, that SOX2 and LGR5 positive supporting cells from the adult organ of Corti are able to respond to Wnt activation, proliferate and differentiate, potentially allowing the use of these compounds to promote hearing restoration in clinical settings.

## Discussion

### Regenerative capacity in the adult mouse cochlea

Cochlear organoids were first developed by McLean et al., 2017, and it refers to primary cochlear progenitor cells grown in 3D as organoids (McLean et al., 2017). They have been mostly produced from neonatal mouse cochlear epithelium (Carpena et al., 2025; Kalra et al., 2023; Kubota et al., 2021; Li & Doetzlhofer, 2020; Liu et al., 2021; Ma et al., 2022; Wu et al., 2025; Xia et al., 2023) mainly because the cochlea is still under development in the early postnatal week in mice leading to enhanced regeneration in neonatal mice accompanied by the epigenetic modifications that hide *Atoh1* transcription factors leading to reduced regeneration in adulthood (Deng & Hu, 2020; Tao et al., 2021). These studies have widely demonstrated that the neonatal cochlea possesses the ability to proliferate, differentiate to Myosin 7A positive cells and even to promote extended passaging in the presence of CHIR99021 and LPA in the expansion medium (Xia et al., 2023). Although adult mammalian cochlear regeneration was believed to be non-existent or dormant, very limited number of publications have shown the differences between early postnatal and adult cochlear regeneration *in vitro*. Oshima et al., 2007 showed that the organoid generating potential is severely reduced from p1 to p21 and onwards in mice, with no spheroid formation after p21 in samples from the organ of Corti (Oshima et al., 2007). While others evaluated adult maculae and sacculae but not organ of Corti regeneration *in vitro* (White et al., 2006). Even though these publications come from a time where Matrigel and Lgr5-GFP animal models had not been developed, they shed insights into a severely reduced mammalian adult regenerative capacity. Another contributor to reduced mammalian adult regeneration is the epigenetic barrier. It has been demonstrated that the epigenetic barrier hampers the regeneration of HCs between postnatal day 7 and 10 in mice (Deng & Hu, 2020; Tao et al., 2021). The modulation of the epigenetic barrier has been used as a mechanism to activate HC regeneration in different models (Layman & Zuo, 2014; McLean et al., 2017; Tang et al., 2016). Although the previously published protocols for the development of cochlear organoids include VPA as HDAC inhibitor to enhance *Atoh1* expression and further activate regeneration (McLean et al., 2017; Xia et al., 2023), in our protocol the activation of the Wnt pathway, via surrogate Wnt, and co-activation of the Lgr5 receptor, via Rspondin1, is sufficient to reactivate these cells to proliferate and differentiate to Myosin 7A positive cells *in vitro* and *ex vivo*.

In order to model ageing in our animal model, we used the old breeder mice used in our breeding protocols (also in line with the 3Rs principles) to evaluate the capacity of the aged cochlea to proliferate and regenerate *in vitro* with our optimized protocol. Our aged model consisted of >p150 C57BL/6J WT and Lgr5-GFP mice (with C57Bl6/J background) that are prone to early onset of hearing loss due to a mutation in the cadherin 23 gene (Cdh23), encoding a component of the HC tip link (Keithley et al., 2004). We confirmed increased ABR thresholds and significantly reduced amplitudes in wave I. We demonstrated that the aged cochlea has the capacity to produce cochlear organoids, and to regenerate *in vitro*. We even observed an enhanced regenerative capacity in the aged cochlea compared to the young adult cochlea, suggesting that the damage that occurs during ageing is priming the cochlea for enhanced *in vitro* regeneration.

### Reduced regeneration capacity in the Lgr5-GFP mouse model

We have evaluated the differences between WT and Lgr5-GFP transgenic mouse derived cochlear progenitors in their capacity to produce cochlear organoids. The Lgr5-GFP transgenic mouse model has been used world-wide for the discovery, study and tracking of the Lgr5 positive cells throughout the organism, including but not limited to intestine, stomach, liver, lungs, mammary and prostate glands, pancreas, cochlea, olfactory system, eye ovaries (Leung et al., 2018). However, the Lgr5-GFP transgenic mouse model has several limitations for the study of non-regenerative organs. Since the EGFP expression is driven by the Lgr5 promoter, one allele has a truncated Lgr5 copy, which suggests that less Lgr5 mRNA is translated. Although this has been shown for the intestine (Tan et al., 2021), it has not been demonstrated for the cochlea. We previously analyzed the *Lgr5* mRNA expression in aged WT and Lgr5-GFP transgenic mice and found no differences (Smith-Cortinez et al., 2021). This does not mean that there might not be differences in younger mice. Because of the clear differences we observed in number and sizes of organoids produced per cochlea in age matched WT versus Lgr5-GFP mice, here we used Lgr5-GFP mice to produce organoids to validate LGR5 expression in expansion and differentiation protocols. However we used WT mice for the rest of the experiments to evaluate the true regenerative potential of the adult cochlea, which is dependent on LGR5. The striking differences between Lgr5-GFP and WT mice in their capacity to produce cochlear organoids further confirms the hypothesis that the LGR5 positive cells are the ones that respond to Wnt signaling to produce Myosin 7A positive cells *in vitro*.

### Regenerative capacity of the cochlea and vestibular organ

It has been suggested that, in mammals, the sensory epithelium of the vestibular organ has increased regenerative capacity compared to the cochlea. Senn et al., 2020 showed that the utricle and ampulla have enhanced success rate in producing spheres than the cochlea in surgical and post mortem derived samples. We compared the proliferative and regenerative potential of the adult mouse and human cochlea to the vestibular organ. We observed the vestibular organ derived samples were able to produce more organoids from human and mouse epithelium; however, they showed similar Myosin 7A expression after differentiation. These findings suggest the presence of more progenitor cells or more cells that have the capacity to respond to Wnt activation. Still, the signaling pathways that drive type I and type II HC in vestibular organ are slightly different than the cochlea, suggesting the protocols might need to be tailored to vestibular organ for adult derived epithelia.

## Conclusions

Here, we have shown for the first time that human adult cochlear and vestibular organ sensory epithelia possess progenitor potential and express LGR5 and SOX2. We showed that the adult and aged human and mouse cochlea possess progenitor potential, express LGR5 and SOX2, and have the capacity to generate organoids *in vitro* using an optimized culture media composition. These organoids differentiate to Myosin 7A positive cells with polarized F-Actin, FM1-43 uptake, stereocilia-like structures, indicative of functional HCs. In conclusion, this protocol can be used for the development of drug testing platforms to test novel drugs to treat hearing loss. Moreover, the compounds used in the optimized medium could potentially promote hearing restoration in clinical settings.

## Author contributions

Conceptualization: NS-C, LVS, GF; Data curation: NS-C, GF, TP; Formal analysis: NS-C, GF, TP; Funding acquisition – NS-C, LVS, RJS; Investigation: : NS-C, GF, TP, CC, SS, FR, CdH, NL, HGXMT; Methodology: NS-C, GF, TP. Project administration: NS-C, LVS, RS; Resources: HGXMT, CC, FR, SS, NL, PS. Supervision: NS-C, LVS; Visualization: GF; Writing – original draft: NS-C; Writing – review & editing: NS-C, LVS, GF, TP, FR, CC

## Supplementary Material

**Supplementary Figure 1.**
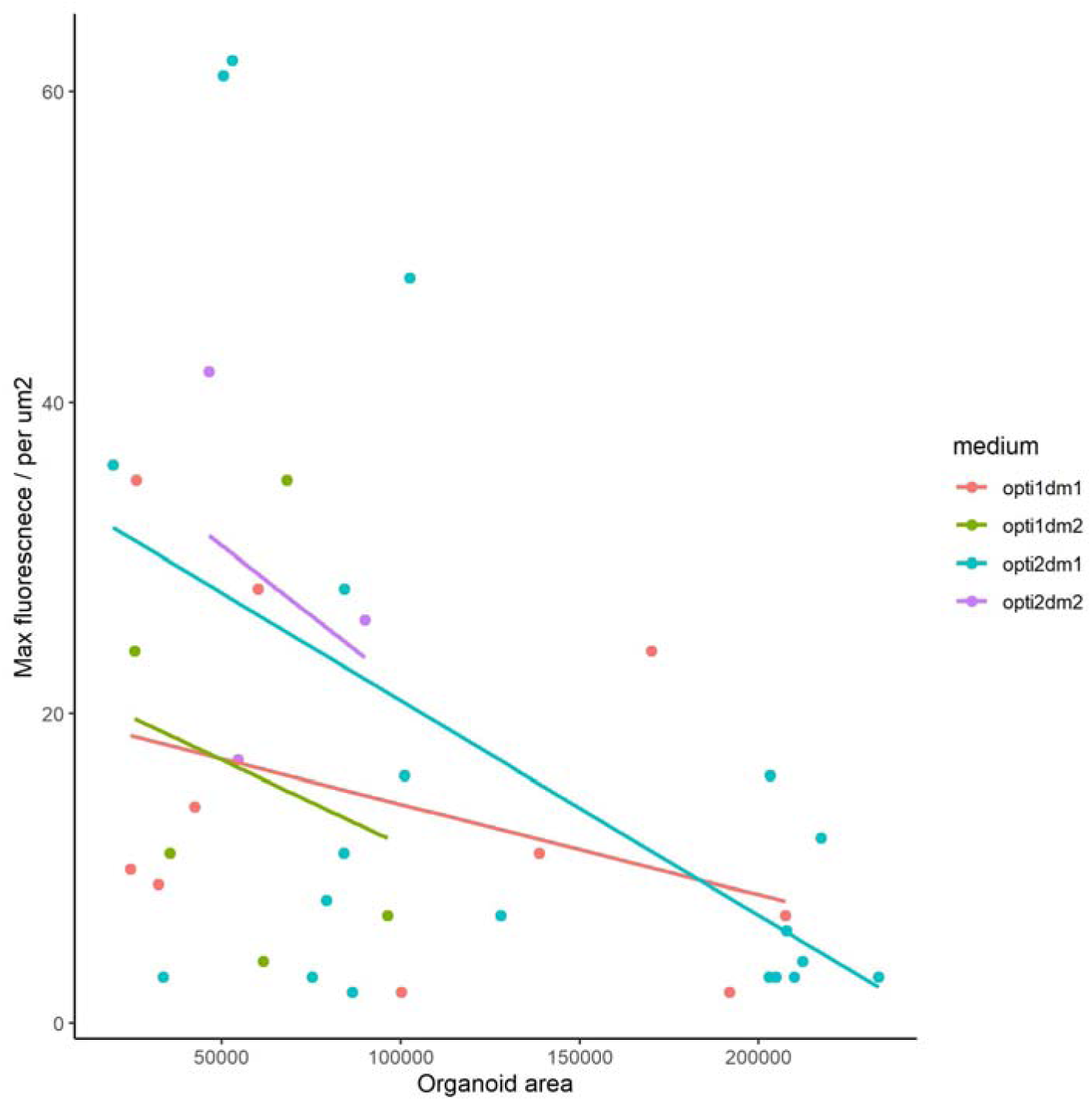
Myosin 7A expression versus size of organoids.

**Supplementary Figure 2:**
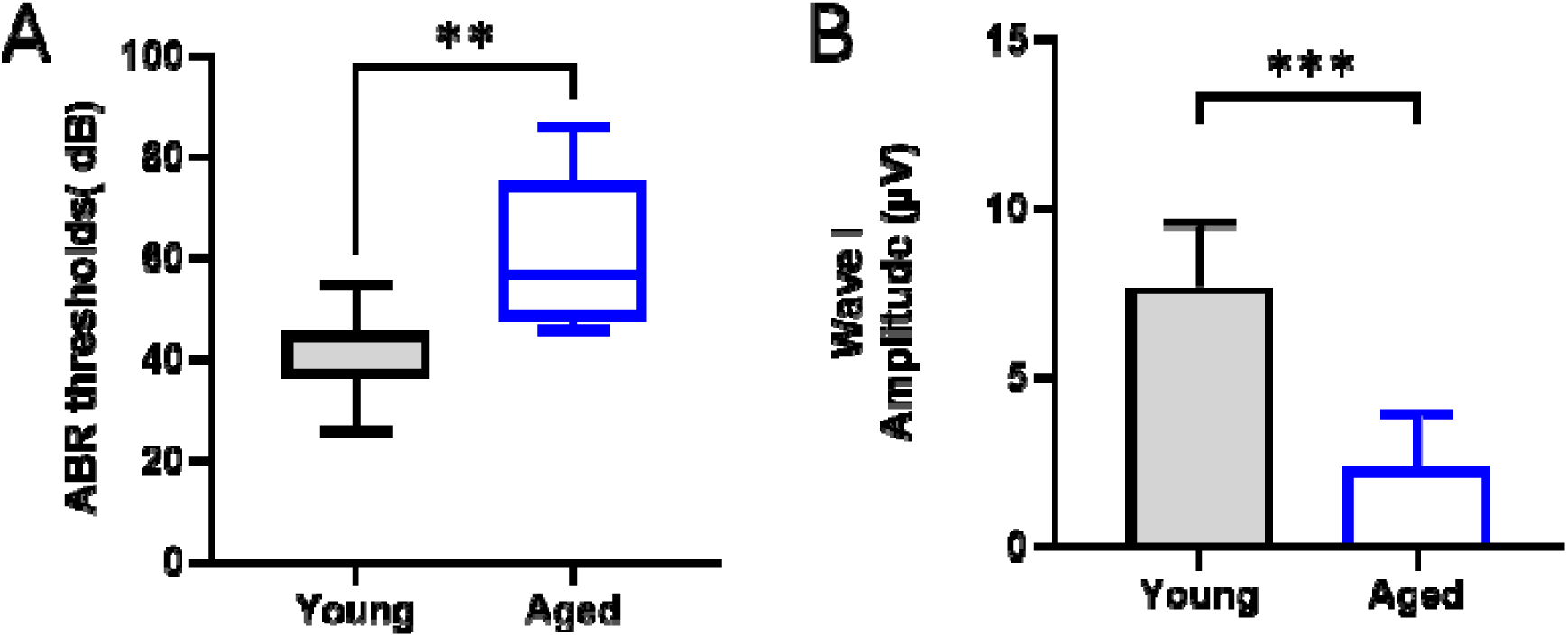
ABR thresholds and Wave I amplitudes in young versus aged mice.

**Supplementary Figure 3:**
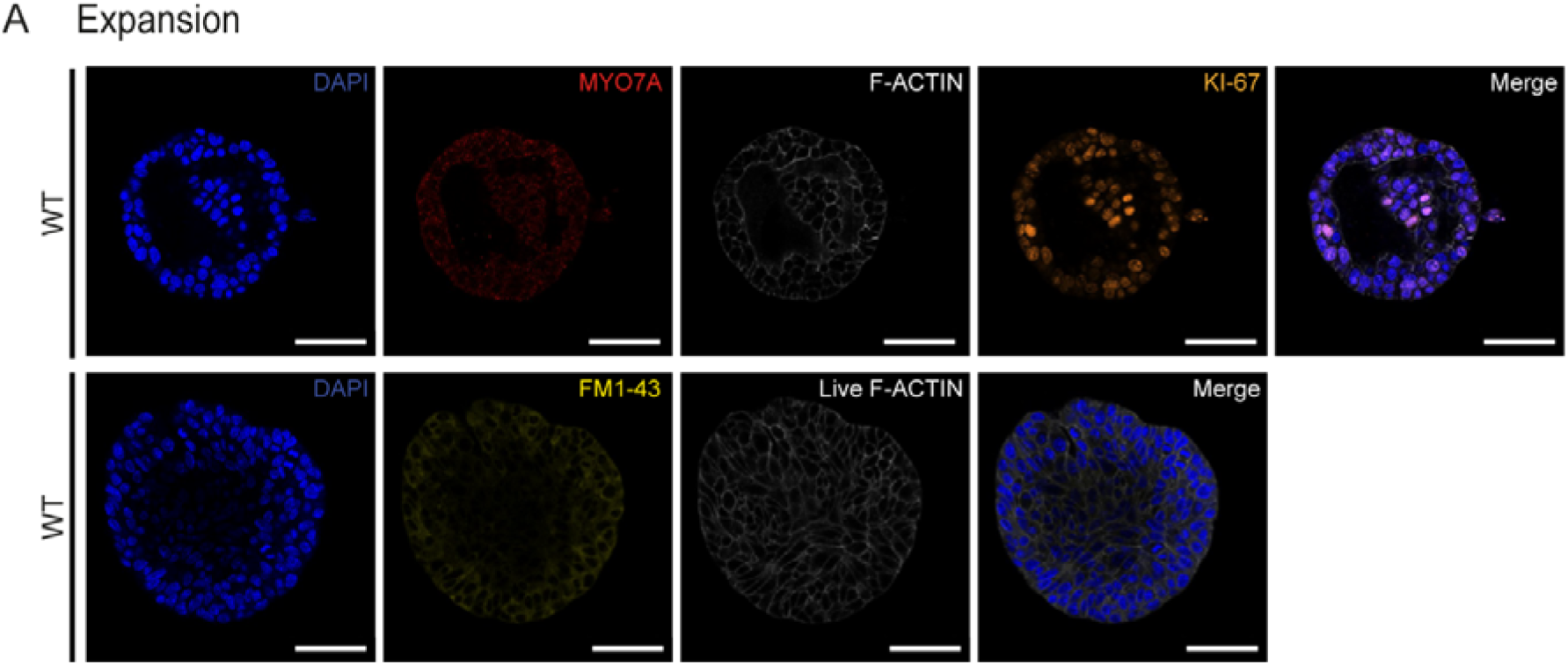
Expression of hair cell markers in cochlear organoids grown in OPTI2 DM medium.

**Supplementary Table 1:** Growth of cochlear organoids in McLean EM on day 15 after isolation for cochlea and vestibular organ; and age of patients.

| Sample # | Age | Number of organoids |  |
| --- | --- | --- | --- |
|  |  | Vestibular organ | Cochlea |
| 1 | 60 | 14 | 15 |
| 2 | 69 | 5 | 0 |
| 3 | 50 | 5 | 2 |
| 4 | 54 | 7 | 0 |
| 5 | 52 | 35 | 0 |
| 6 | 48 | 20 | 6 |
| 7 | 35 | 21 | 5 |
| 8 | 23 | 1 | 0 |
| 9 | 43 | 3 | 4 |
| 10 | 59 | 6 | 8 |

**Supplementary Table 2:** Characteristics of body donors included in study.

| Sample # | Sex | Age | Post mortem interval (hrs) |
| --- | --- | --- | --- |
| 1 | M | 65 | 14 |
| 2 | F | 79 | 25 |
| 3 | M | 80 | 31 |
| 4 | M | 87 | 10 |
| 5 | M | 87 | 8-10 |

M: Male, F: Female

**Supplementary Table 3:**
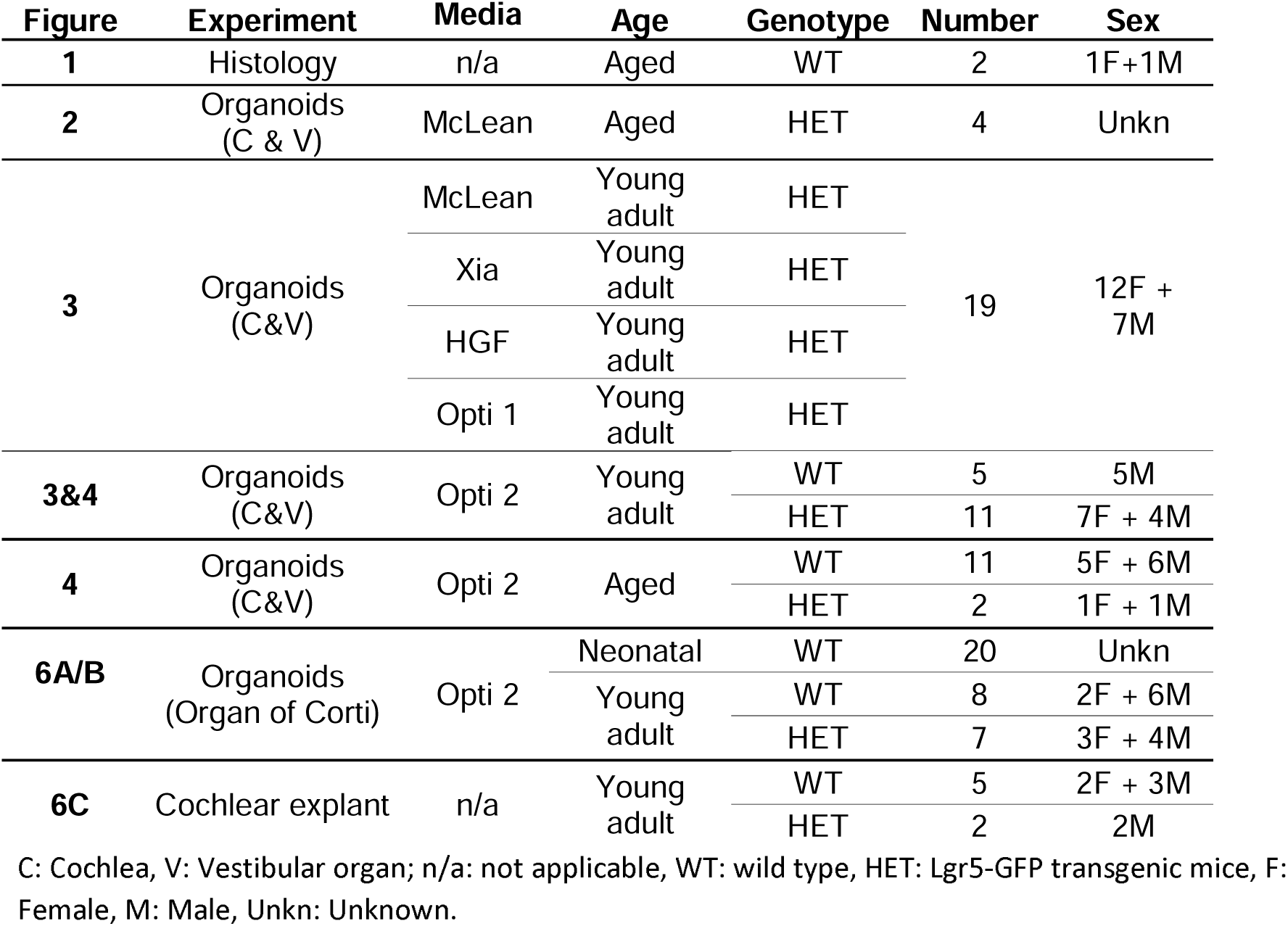
Overview of all animals used in each Figure, Experiment, Aged.

| Figure | Experiment | Media | Age | Genotype | Number | Sex |
| --- | --- | --- | --- | --- | --- | --- |
| **1** | Histology | n/a | Aged | WT | 2 | 1F+1M |
| **2** | Organoids (C & V) | McLean | Aged | HET | 4 | Unkn |
| **3** | Organoids (C&V) | McLean | Young adult | HET | 19 | 12F + 7M |
|  |  | Xia | Young adult | HET |  |  |
|  |  | HGF | Young adult | HET |  |  |
|  |  | Opti 1 | Young adult | HET |  |  |
| **3&4** | Organoids (C&V) | Opti 2 | Young adult | WT | 5 | 5M |
|  |  |  |  | HET | 11 | 7F + 4M |
| **4** | Organoids (C&V) | Opti 2 | Aged | WT | 11 | 5F + 6M |
|  |  |  |  | HET | 2 | 1F + 1M |
| **6A/B** | Organoids (Organ of Corti) | Opti 2 | Neonatal | WT | 20 | Unkn |
|  |  |  | Young adult | WT | 8 | 2F + 6M |
|  |  |  |  | HET | 7 | 3F + 4M |
| **6C** | Cochlear explant | n/a | Young adult | WT | 5 | 2F + 3M |
|  |  |  |  | HET | 2 | 2M |

C: Cochlea, V: Vestibular organ; n/a: not applicable, WT: wild type, HET: Lgr5-GFP transgenic mice, F: Female, M: Male, Unkn: Unknown.

**Supplementary Table 4:** High Growth Factor Expansion Medium (HGF)

| Compound | Description | Stock Concentration | Working Concentration |
| --- | --- | --- | --- |
| DMEM/F12 | Basal medium | N/A | N/A |
| WNT-CM* | Wnt-CM* | N/A | N/A |
| B27 Supplement 50x | Growth factor supplement | N/A | N/A |
| N2 Supplement 100x | Growth factor supplement | N/A | N/A |
| Ascorbic Acid | Small molecule | 100 mg/mL | 100 µg/mL |
| EGF | Growth factor | 100 µg/mL | 50 ng/mL |
| bFGF | Growth factor | 100 µg/mL | 50 ng/mL |
| IGF-10 | Growth factor | 100 µg/mL | 50 ng/mL |
| RepSox | Small molecule | 20 mM | 2 µM |
| R-Spondin | Ligand | 250 µg/mL | 500 ng/mL |
| Noggin | Ligand | 100 µg/mL | 10 µg/mL |

\* Wnt-CM: Wnt conditioned medium

**Supplementary Table 5:** Optimized isolation medium (Opti IM)

| Compound | Description | Stock Concentration | Working Concentration |
| --- | --- | --- | --- |
| DMEM/F12 | Basal medium | N/A | N/A |
| B27 Supplement 50x | Growth factor supplement | N/A | N/A |
| N2 Supplement 100x | Growth factor supplement | N/A | N/A |
| NAC | Antioxidant | 125 mM | 1.25 mM |
| Ascorbic Acid | Small molecule | 100 mg/mL | 100 µg/mL |
| EGF | Growth factor | 100 µg/mL | 50 ng/mL |
| bFGF | Growth factor | 100 µg/mL | 50 ng/mL |
| IGF-10 | Growth factor | 100 µg/mL | 50 ng/mL |
| Surrogate Wnt | Small molecule | 1 mM | 1 nM |
| RepSox | Small molecule | 20 mM | 2 µM |
| R-Spondin | Ligand | 250 µg/mL | 500 ng/mL |
| Y-27632 | Ligand | 10 mM | 10 µM |

**Supplementary Table 6:** Optimized expansion medium 1 (Opti1 EM)

| Compound | Description | Stock Concentration | Working Concentration |
| --- | --- | --- | --- |
| DMEM/F12 | Basal medium | N/A | N/A |
| B27 Supplement 50x | Growth factor supplement | N/A | N/A |
| N2 Supplement 100x | Growth factor supplement | N/A | N/A |
| NAC | Antioxidant | 125 mM | 1.25 mM |
| Ascorbic Acid | Small molecule | 100 mg/mL | 100 µg/mL |
| EGF | Growth factor | 100 µg/mL | 50 ng/mL |
| bFGF | Growth factor | 100 µg/mL | 50 ng/mL |
| IGF-10 | Growth factor | 100 µg/mL | 50 ng/mL |
| CHIR99021 | Small molecule | 25 mM | 3 µM |
| RepSox | Small molecule | 20 mM | 2 µM |
| R-Spondin | Ligand | 250 µg/mL | 500 ng/mL |
| LPA | Small molecule | 1 mM | 2.5 µM |

**Supplementary Table 7:** Optimized expansion medium 2 (Opti2 EM)

| Compound | Description | Stock Concentration | Working Concentration |
| --- | --- | --- | --- |
| DMEM/F12 | Basal medium | N/A | N/A |
| B27 Supplement 50x | Growth factor supplement | N/A | N/A |
| N2 Supplement 100x | Growth factor supplement | N/A | N/A |
| NAC | Antioxidant | 125 mM | 1.25 mM |
| Ascorbic Acid | Small molecule | 100 mg/mL | 100 µg/mL |
| EGF | Growth factor | 100 µg/mL | 50 ng/mL |
| bFGF | Growth factor | 100 µg/mL | 50 ng/mL |
| IGF-10 | Growth factor | 100 µg/mL | 50 ng/mL |
| Surrogate Wnt | Small molecule | 1 mM | 1 nM |
| RepSox | Small molecule | 20 mM | 2 µM |
| R-Spondin | Ligand | 250 µg/mL | 500 ng/mL |
| LPA | Small molecule | 1 mM | 2.5 µM |

**Supplementary Table 8:** Optimized differentiation medium 1 (Opti DM1)

| Compound | Description | Stock Concentration | Working Concentration |
| --- | --- | --- | --- |
| DMEM/F12 | Basal medium | N/A | N/A |
| B27 Supplement 50x | Growth factor supplement | N/A | N/A |
| N2 Supplement 100x | Growth factor supplement | N/A | N/A |
| NAC | Antioxidant | 125 mM | 1.25 mM |
| Ascorbic Acid | Small molecule | 100 mg/mL | 100 µg/mL |
| Shh | Small molecule | 25 µg/mL | 100 ng/mL |
| Surrogate Wnt | Small molecule | 1 mM | 1 nM |
| RepSox | Small molecule | 20 mM | 2 µM |
| LY411575 | Small molecule | 10 mM | 5 µM |

**Supplementary Table 9:**
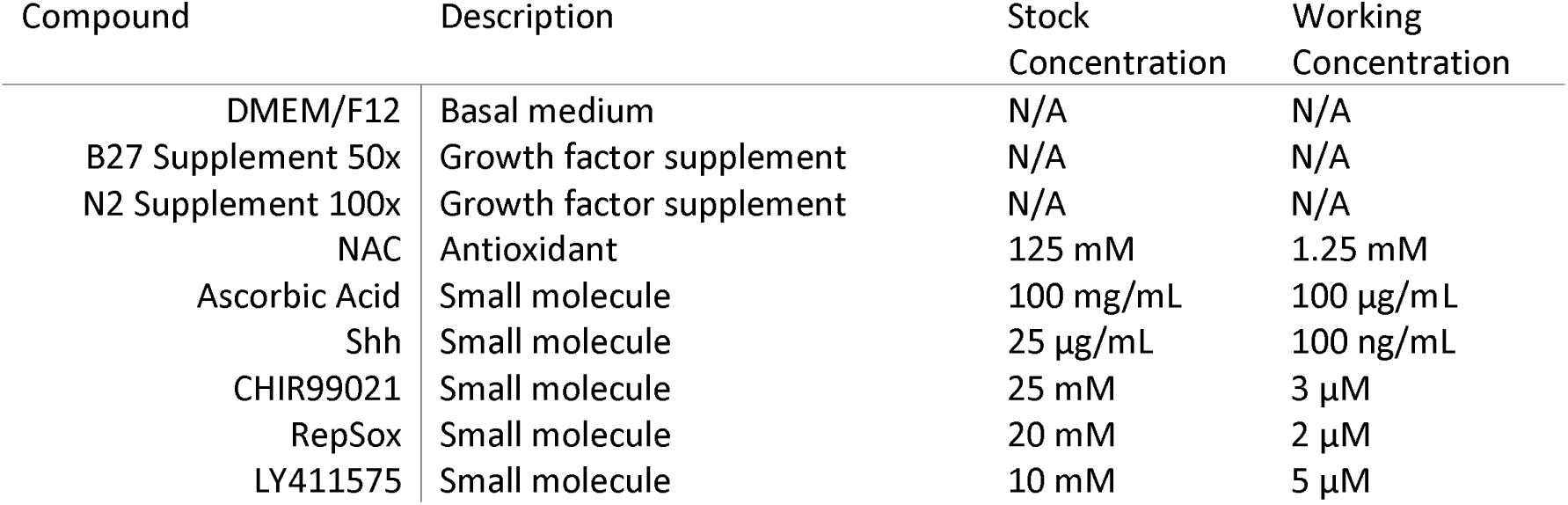
Optimized differentiation medium 2 (Opti DM2)

| Compound | Description | Stock Concentration | Working Concentration |
| --- | --- | --- | --- |
| DMEM/F12 | Basal medium | N/A | N/A |
| B27 Supplement 50x | Growth factor supplement | N/A | N/A |
| N2 Supplement 100x | Growth factor supplement | N/A | N/A |
| NAC | Antioxidant | 125 mM | 1.25 mM |
| Ascorbic Acid | Small molecule | 100 mg/mL | 100 µg/mL |
| Shh | Small molecule | 25 µg/mL | 100 ng/mL |
| CHIR99021 | Small molecule | 25 mM | 3 µM |
| RepSox | Small molecule | 20 mM | 2 µM |
| LY411575 | Small molecule | 10 mM | 5 µM |

**Supplementary Table 10:** ABR thresholds and Wave I amplitudes in young adult versus aged C57BL/6J and Lgr5-GFP mice.

| Mouse # | Age (Y/A) | Genotype | Sex | Threshold (Db attenuation) | ABR Threshold | Wave I amplitude (μV) |
| --- | --- | --- | --- | --- | --- | --- |
| M053 | Y | Het | M | 60 | 45 | 7.5 |
| M054 | Y | Het | M | 60 | 45 | 8.7 |
| M055 | Y | Het | M | 61 | 44 | 3.9 |
| M582 | Y | Het | M | 69 | 36 | 7.6 |
| M583 | Y | WT | M | 59 | 46 | 8.8 |
| M584 | Y | WT | M | 68 | 37 | 7.1 |
| M585 | Y | WT | F | 71 | 34 | 6.5 |
| M588 | Y | WT | F | 79 | 26 | 10.6 |
| M589 | Y | WT | F | 56 | 49 | 9.2 |
| M593 | Y | WT | M | 63 | 42 | 8.1 |
| M595 | Y | WT | M | 61 | 44 | 4.4 |
| M596 | Y | WT | F | 60 | 45 | 9.8 |
| M597 | Y | Het | F | 60 | 45 | 7.3 |
| M032 | A | WT | M | 57 | 48 | 3.1 |
| M033 | A | WT | M | 48 | 57 | 4.3 |
| M204 | A | WT | M | 35 | 70 | 1.7 |
| M207 | A | WT | M | 59 | 46 | 4 |
| M208 | A | WT | M | 53 | 52 | 4.2 |
| M443 | A | WT | M | 47 | 58 | 1.6 |
| M193 | A | Het | F | 25 | 80 | 1.7 |
| M618 | A | WT | F | 19 | 86 | 0.3 |
| M346 | A | Het | M | 57 | 48 | 3.9 |

